# Recombinant outer membrane vesicles coupled with cytokines act as high-performance adjuvants against *Helicobacter pylori* infection in mice

**DOI:** 10.1101/2023.06.26.546595

**Authors:** Qiong Liu, Biaoxian Li, Jiahui Lu, Yejia Zhang, Yinpan Shang, Yi Li, Tian Gong, Chengsheng Zhang

## Abstract

The widespread prevalence of *Helicobacter pylori* (*H. pylori*) infection remains a great challenge to the human health. The existing vaccines are not ideal for preventing *H. pylori* infection; thus, exploring highly effective adjuvants may improve the immunoprotective efficacy of *H. pylori* vaccines. In a previous study, we found that outer membrane vesicles (OMVs), a type of nanoscale particle spontaneously produced by Gram-negative bacteria, could act as adjuvants to boost the immune response to vaccine antigens. In the present study, we explored the potential application of OMVs as delivery vectors for adjuvant development. We constructed recombinant OMVs containing cytokines interleukin 17A or interferon-γ and evaluated their function as adjuvants in combination with inactivated whole-cell vaccine (WCV) or UreB as vaccine antigens. Our results showed that recombinant OMVs as adjuvants could induce stronger humoral and mucosal immune responses in mice than wild-type *H. pylori* OMVs and the cholera toxin (CT) adjuvant. Additionally, the recombinant OMVs significantly promoted Th1/Th2/Th17-type immune responses. Furthermore, the recombinant OMV adjuvant induced more potent clearance of *H. pylori* than CT and wild-type OMVs. Our data suggest that the recombinant OMVs coupled with cytokines may become potent adjuvants for development of novel and effective vaccines against *H. pylori* infection.

**Importance:** *Helicobacter pylori* (*H. pylori*) is one of the important risk factors for gastric cancer, and its vaccine is crucial for its prevention and control. However, to date, no effective vaccine has been approved. Exploring novel and effective vaccine adjuvants may provide new perspectives and ideas for the development of *H. pylori* vaccines. Outer membrane vesicles (OMVs) deserve more attention as a novel form of vaccine and adjuvant. We have long focused on OMVs as vaccine adjuvants to enhance efficacy by delivering eukaryotic plasmid expressing immune promoting cytokines in wild-type OMVs. Our results are expected to provide a new adjuvant form for the development of *H. pylori* vaccine, and this adjuvant design strategy can also be used in the development of vaccines for other types of pathogens, including bacteria and even viruses.

## 1. Introduction

*Helicobacter pylori* (*H. pylori*), the most important risk factor for gastric cancer, causes a large number of infections worldwide and poses a great danger to human health; thus, the early prevention of *H. pylori* infection needs urgent attention^1,2^. To find effective treatments for *H. pylori* infection, many studies have focused on the development of *H. pylori* vaccines based on inactivated whole bacteria and immunoprotective proteins or peptides^3^. Numerous studies have also screened for antigenic components with good protective efficacy for vaccine design, such as various antigenic epitopes against UreB proteins^4,5^. However, neither whole-cell inactivated vaccines alone nor the antigenic proteins or highly potent epitopes induce an immune response strong enough to clear *H. pylori* infection^6^. The development of novel vaccine adjuvants that can improve their protective efficacy may be the best strategy to address this issue and needs to be a priority in *H. pylori* vaccine development.

The ideal *H. pylori* vaccine adjuvant should be safe, promote a durable, high-quality immune response to the antigen, and induce a mucosal immune response and a T helper (Th)1/Th17-type immune response that significantly enhances the capacity of the host to prevent *H. pylori* infection^7–9^. Currently, the adjuvants used in *H. pylori* whole-cell bacterial vaccines or recombinant *H. pylori* vaccines include aluminum adjuvant, cholera toxin, heat-insensitive *E. coli* enterotoxin, complete and incomplete Freund’s adjuvant, CpG oligonucleotides, and chitosan^10,11^. Although these adjuvants have been used in experimental animal models to improve the immune response induced by *H. pylori* vaccine antigen stimulation, all of them have certain drawbacks that affect their clinical application. Therefore, there is a continued need to explore novel vaccine adjuvants to develop *H. pylori* vaccines.

Outer membrane vesicles (OMVs), nanoscale vesicle-like structures spontaneously produced by bacteria, contain multiple pathogen-associated molecular patterns (PAMPs) and exist in a non-replicating form. These characteristics endow them with powerful immunostimulatory ability^12–14^. Previously, most studies focused on the use of OMVs as antigenic components for vaccine development and often overlooked their potential as vaccine adjuvants. The special nanoscale structure of OMVs also allows them to serve as efficient delivery vectors that can be genetically engineered for use as vaccine adjuvants^15,16^. In a previous study, we explored the potential of wild-type *H. pylori* OMVs as adjuvants for vaccine development. Wild-type OMVs were shown to be effective as adjuvants to improve the clearance of *H. pylori* infection. However, in that work, the full potential of OMVs as delivery vectors for adjuvant development was not realized^17,18^. Therefore, in this study, we modified OMVs via genetic engineering to deliver cytokines capable of enhancing adjuvant efficacy, constructed a new generation of OMV adjuvants for *H. pylori* vaccine development, and evaluated their immune mechanisms.

## 2 Results

### 2.1 Construction and identification of eukaryotic expression vectors

First, we constructed eukaryotic expression vectors of cytokines interleukin 17A (IL-17A) or interferon-γ (IFN-γ) linked to the enhanced green fluorescent protein (EGFP) or mCherry fluorescent protein. The expression plasmids were subsequently transferred into *H. pylori* OMVs by electroporation to obtain recombinant OMVs (Figure 1A). To assess the delivery efficiency of OMVs delivering eukaryotic plasmids, we used a laser scanning confocal microscope to examine the expression of intracellular fluorescence, reflecting the ability of OMVs to enter the cells and their distribution after entry, and the expression levels of intracellular IFN-γ and IL-17A, as shown in Figure 1B. The data showed that eukaryotic expression plasmids expressing cytokines IL-17A and IFN-γ were successfully delivered into cells by OMVs, and the incubation of OMVs delivering both cytokines allows these cytokines to enter cells simultaneously. In addition, high levels of cytokines IFN-γ and IL-17A were detected in HEK-293T and GES-1 cells by reverse transcription quantitative PCR (RT-qPCR) and Enzyme-linked immunosorbent assay (ELISA). (Figure 1C, 1D). The above results indicate that plasmids expressing cytokines IFN-γ (EGFP) or IL-17A (mCherry) were successfully delivered into eukaryotic cells by OMVs.

**Figure 1.**
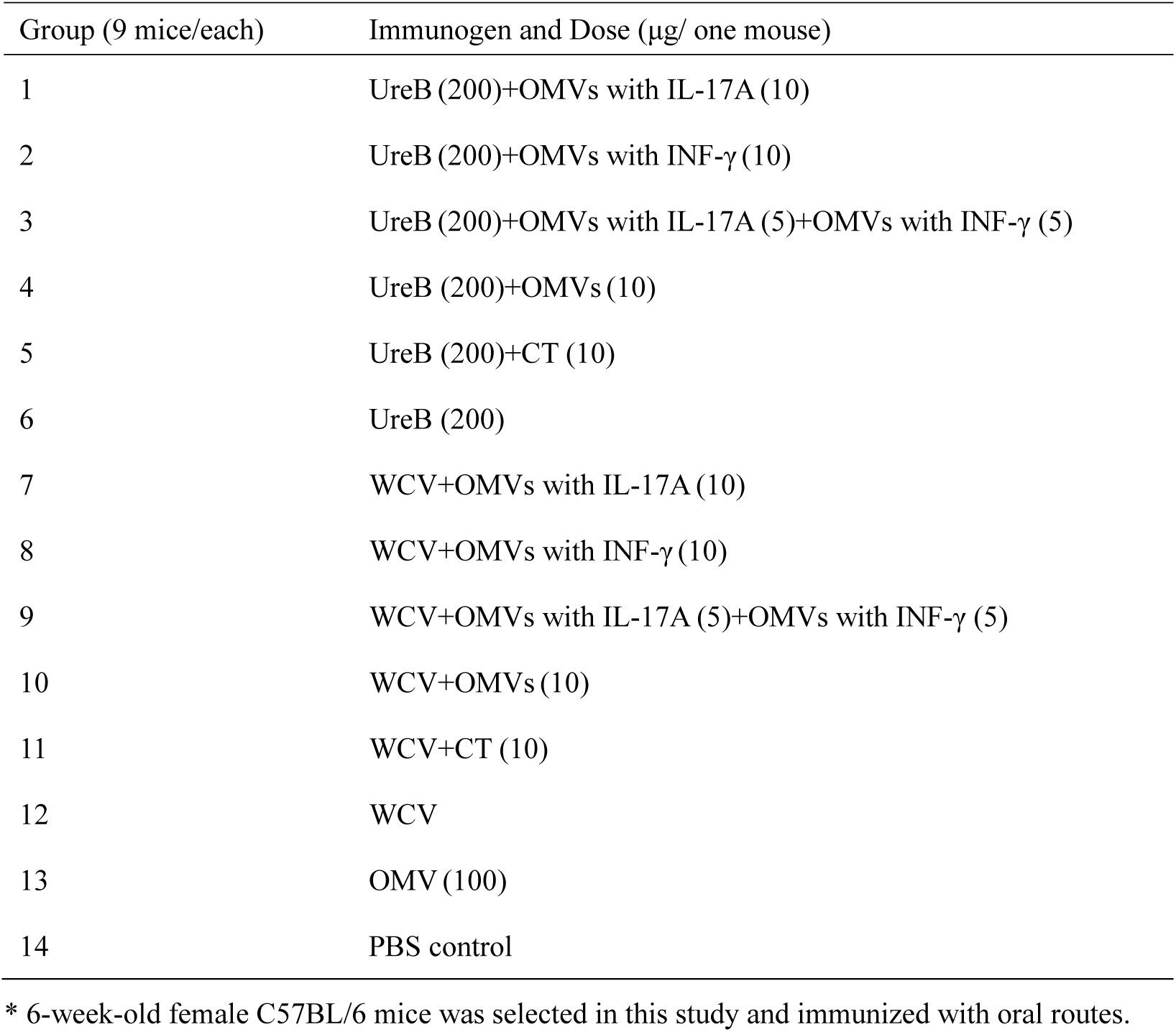
Construction and identification of recombinant OMVs. Construction of recombinant cytokine OMVs and their pattern diagram for immunization as adjuvant (A). HEK-293T and GES-1 cells were divided into three groups each: 100 μg of OMVs delivering cytokine IFN-γ (EGFP) was added to the first group, 100 μg of OMVs delivering cytokine IL-17A (mCherry) was added to the second, and 50 μg each of OMVs delivering cytokine IFN-γ (EGFP) or IL-17A (mCherry) was added to the third, followed by incubation in a cell culture incubator. Eukaryotic expression plasmids are highly expressed in both HEK-293T and GES-1 cells as observed by laser confocal microscopy (B). The expression of IL-17A and IFN-γ in cells by RT-qPCR (C) or quantitative ELISA (D).

### 2.2 Potent systemic immune responses induced by recombinant OMV adjuvants with subunit or inactivated vaccines

Next, we evaluated the levels of the immune response induced by recombinant OMVs as adjuvants using two different vaccine regimens (UreB and whole inactivated cell vaccine, WCV). Our results showed that the level of anti-*H. pylori* IgG antibodies, representing humoral immunity, was significantly higher after the addition of OMVs as adjuvants, regardless of whether UreB or WCV was used as the vaccine antigen. The addition of recombinant IL-17A-OMVs and recombinant IFN-γ-OMVs + recombinant IL-17A-OMVs as adjuvant groups stimulated a more significant and durable immune response compared to the cholera toxin (CT) adjuvant or wild-type *H. pylori* OMVs as the adjuvant (Figure 2B, 2C).

**Figure 2.**
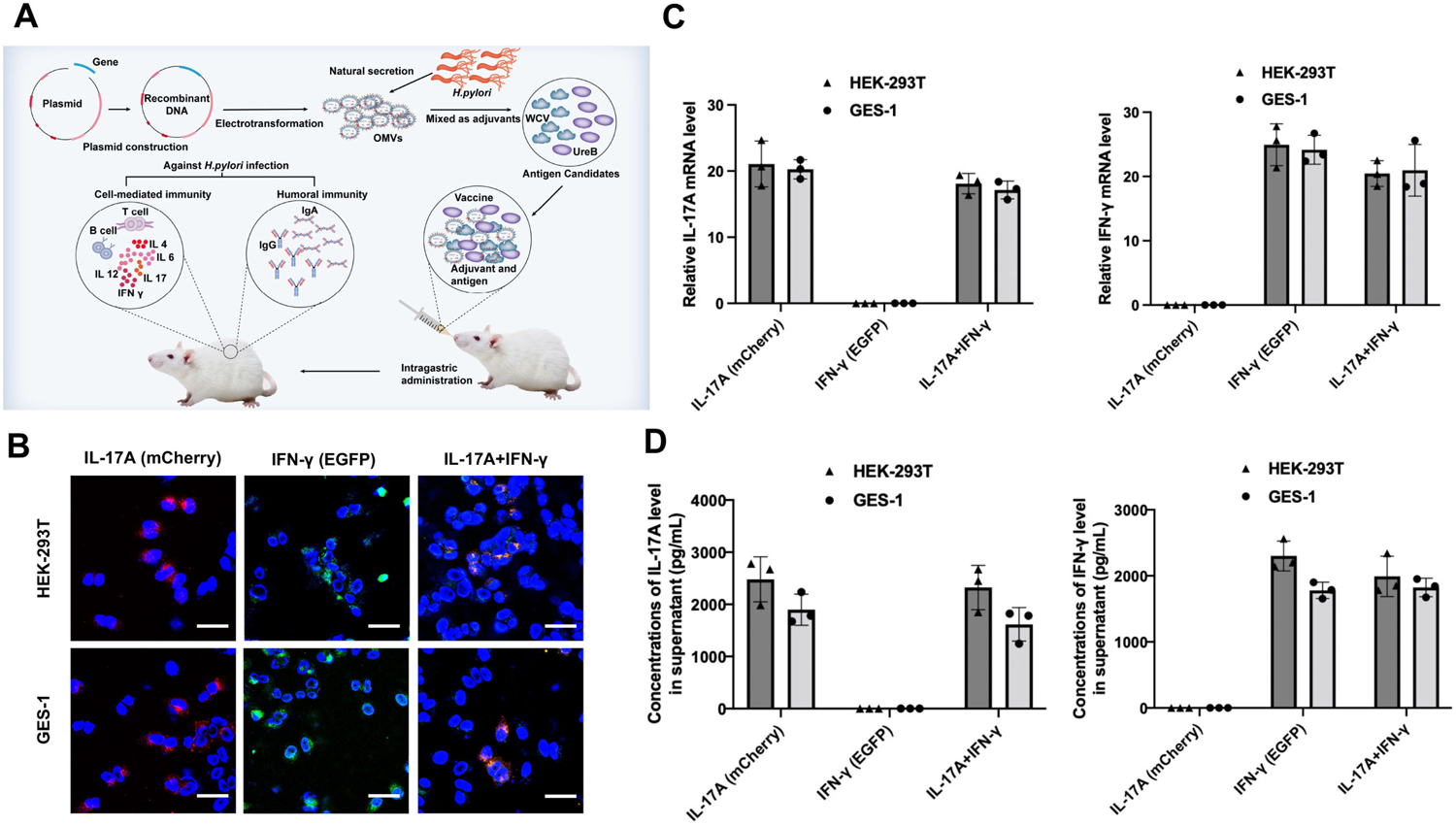
System immune response effects of recombinant OMVs as adjuvants against different types of antigens. Timeline of immunization for this study (A). Serum was collected from mice the day before the first immunization and 2, 4, 6, 8, 10, and 12 weeks after immunization, and serum levels of specific IgG antibodies against UreB and OMP were determined by quantitative ELISA. Wild-type *H. pylori* OMVs and CT were used as control adjuvants, and the PBS group was used as a blank control. Quantitative ELISA to determine the level of anti-*HP* UreB in serum during 12 weeks of immunization in each group when UreB was used as an antigen (B) and the level of anti-*Hp* OMP when WCV was used as an antigen (C). Statistical significance was assessed by two-way ANOVA. *P* < 0.05 and *P* < 0.01 were considered statistically significant. All the results are expressed as mean ± SD per cohort.

We also measured the secretory IgA (S-IgA) in vaginal secretions, the most important indicator of systemic mucosal immunity acquired by vaccination^19^. We observed a phenomenon similar to that seen for IgG. The levels of secretory IgA antibodies produced by the addition of cytokine OMVs as the adjuvant were all significantly higher than those produced in the CT adjuvant group or the wild-type OMV adjuvant group (Supplemental Figure 1A, 1B).

### 2.3 Recombinant OMVs as adjuvants enhance gastric mucosal immune response

Gastric mucosal immunity is the most important line of host defense against *H. pylori* infection, with gastric mucosal secretory IgA directly reflecting the mucosal immune level against *H. pylori* infection^20^., Four mice were sacrificed at week 8 after the first immunization, and their stomachs were taken out aseptically to measure the gastric mucosal IgA content by quantitative ELISA. Our results revealed that when UreB was used as the vaccine antigen, using either recombinant IL-17A-OMVs or recombinant IFN-γ-OMVs as the adjuvant or a combination of recombinant IFN-γ-OMVs + recombinant IL-17A-OMVs as the adjuvant resulted in significantly higher levels of gastric mucosal IgA antibodies than the use of wild-type *H. pylori* OMVs or CT adjuvant (*P* < 0.01; Figure 3A). We obtained similar results when WCV was used as a vaccine antigen, with recombinant OMVs as adjuvants inducing significantly higher levels of stomach IgA antibodies, and recombinant IL-17A-OMVs alone and combined recombinant IFN-γ-OMVs + recombinant IL-17A-OMVs as adjuvants significantly differing from wild-type *H. pylori* OMVs as the adjuvant (Figure 3B).

**Figure 3.**
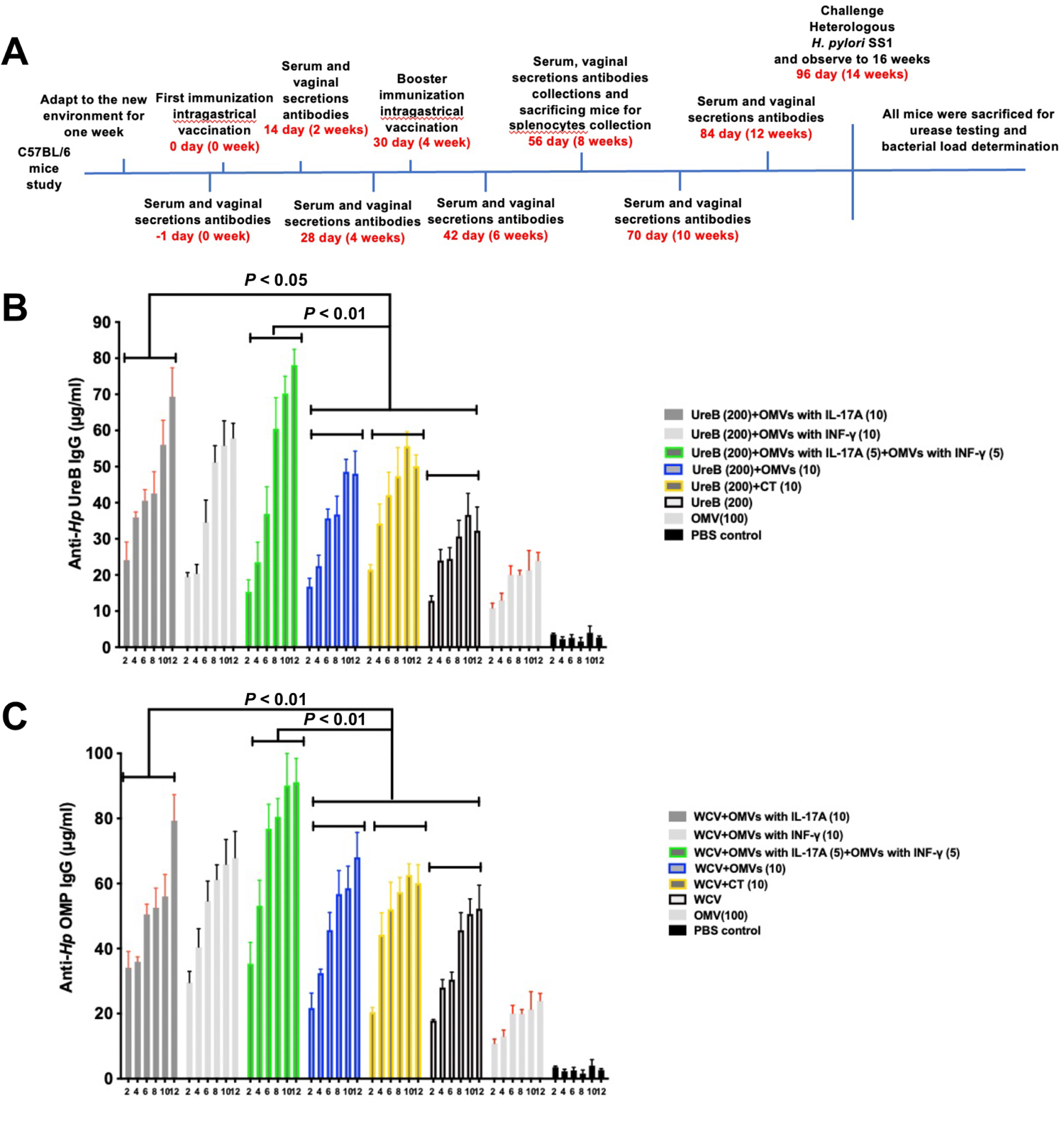
Recombinant OMVs as adjuvants enhance the gastric mucosal immune response. Concentrations of anti-*Hp* UreB stomach IgA using UreB as the vaccine (A) and anti-*Hp* OMP stomach IgA using WCV (B) from the stomach of sacrificed mice were determined by quantitative ELISA in week 8. Each group comprised nine mice. These data displayed the strength of the mucosal immune response induced by diverse immunogens, which are expressed as mean ± SD per group. The least significant difference test was performed to determine whether the distinctions between the means of groups were significant. *P* < 0.05 and *P* < 0.01 represent the differences between the related groups.

### 2.4 Recombinant OMVs as adjuvants strengthen both Th1 and Th2 responses

Next, we examined the expression levels of two subtypes of serum antibody IgG (i.e. IgG_1_and IgG_2c_), produced after immunization with each group. The secretion levels of the IgG_1_ and IgG_2c_ isoforms can reflect the type of immune response generated, which will help understand the molecular mechanism by which recombinant OMVs act as adjuvants to confer immune protection. As seen in Figure 4, the use of recombinant OMVs as adjuvants could stimulate the production of higher levels of IgG_1_ and IgG_2c_ than the CT adjuvant. The combination of recombinant IFN-γ-OMVs + recombinant IL-17A-OMVs produced higher levels of IgG_1_ antibodies compared to recombinant IL-17A-OMVs or recombinant IFN-γ-OMVs alone (*P* < 0.05; Figure 4A, 4C). Regarding recombinant IL-17A-OMVs or recombinant IFN-γ-OMVs alone and recombinant IL-17A-OMVs + recombinant IFN-γ-OMVs in combination, the levels of IgG_2c_ produced by the three groups differed significantly from those produced by wild-type *H. pylori* OMVs as the adjuvant, but there was little difference between the three groups (Figure 4B, 4D). In mice, higher levels of IgG_1_ antibodies represent a Th2-type immune response, while higher levels of IgG_2c_ antibodies reflect a Th1-type immune response^21^. Collectively, the results suggest that recombinant OMVs as adjuvants can induce higher levels of a Th1-type immune response.

**Figure 4.**
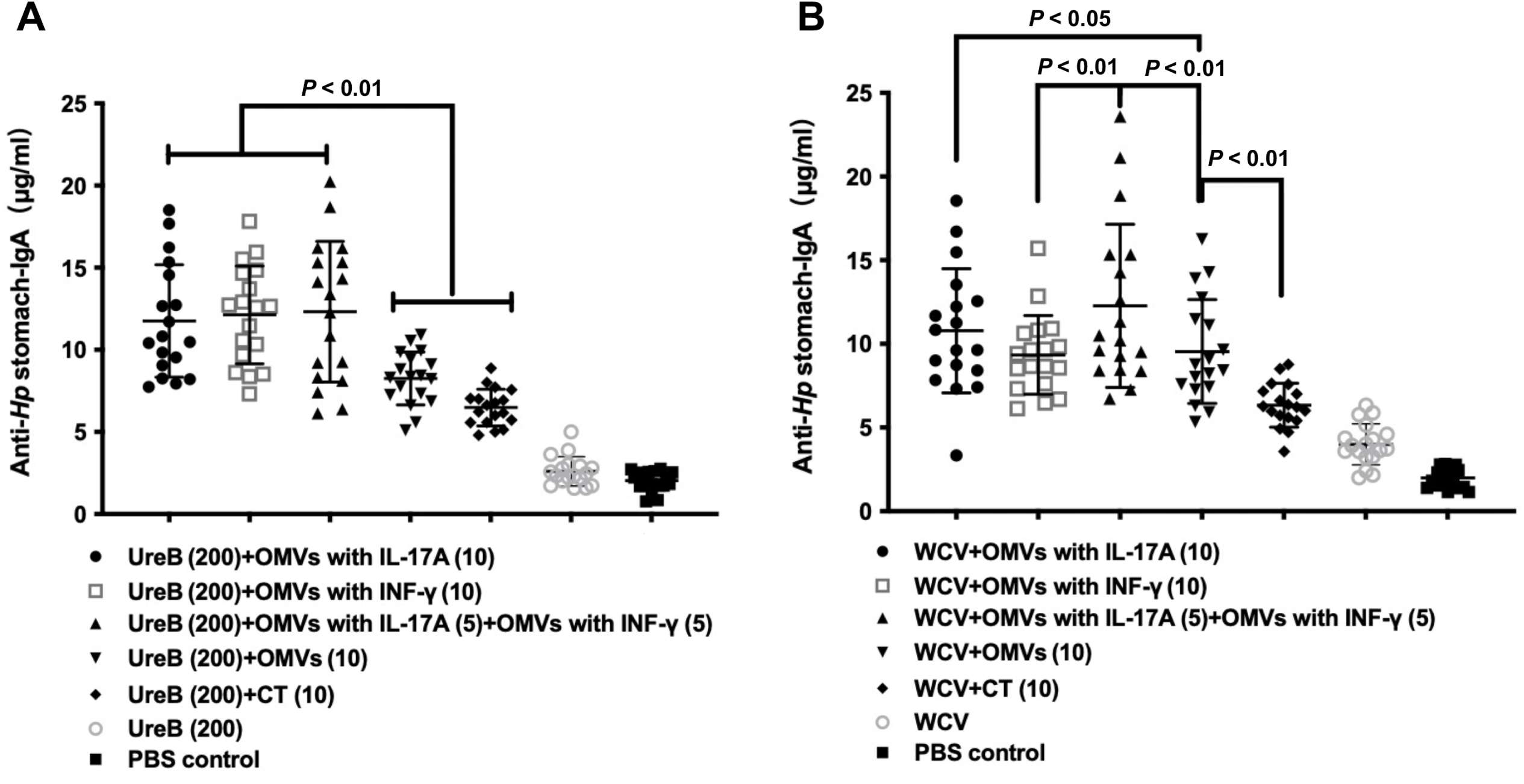
Recombinant OMVs as adjuvants strengthen both Th1 and Th2 responses. The concentrations of anti-*Hp* UreB IgG_1_ (A) and anti-*Hp* OMP IgG_1_ (C) in groups immunized with UreB or WCV as vaccine antigens and the levels of anti-*Hp* UreB IgG_2c_ (B) and anti-*Hp* OMP IgG_2c_ (D) in groups with UreB or WCV were determined by quantitative ELISA on UreB and OMPs isolated from *H. pylori*. Each group consisted of nine mice. The exact concentrations of IgG_1_ and IgG_2c_ subclass antibodies in serum samples from mice 8 weeks after immunization are shown. Means were compared using the least significant difference test. *P* < 0.05 and *P* < 0.01 reflect statistically significant differences between the groups of interest.

### 2.5 Recombinant OMVs as adjuvants induce a stronger Th1/Th2/Th17 balanced immune response

We isolated monocytes from mesenteric lymph node (MLN) and the spleen and examined the secreted cytokine levels to determine the type of immune response enhanced by recombinant OMV adjuvants. In the present study, we first assessed the levels of cytokines IL-17 and IFN-γ in each group after 8 weeks of immunization. Figure 5 clearly shows that the combination of recombinant IFN-γ-OMVs + recombinant IL-17A-OMVs as the adjuvant could significantly increase the levels of cytokines IL-17 and IFN-γ secreted by MLN cells and monocytes compared to wild-type *H. pylori* OMVs as the adjuvant, regardless of the antigen used.

**Figure 5.**
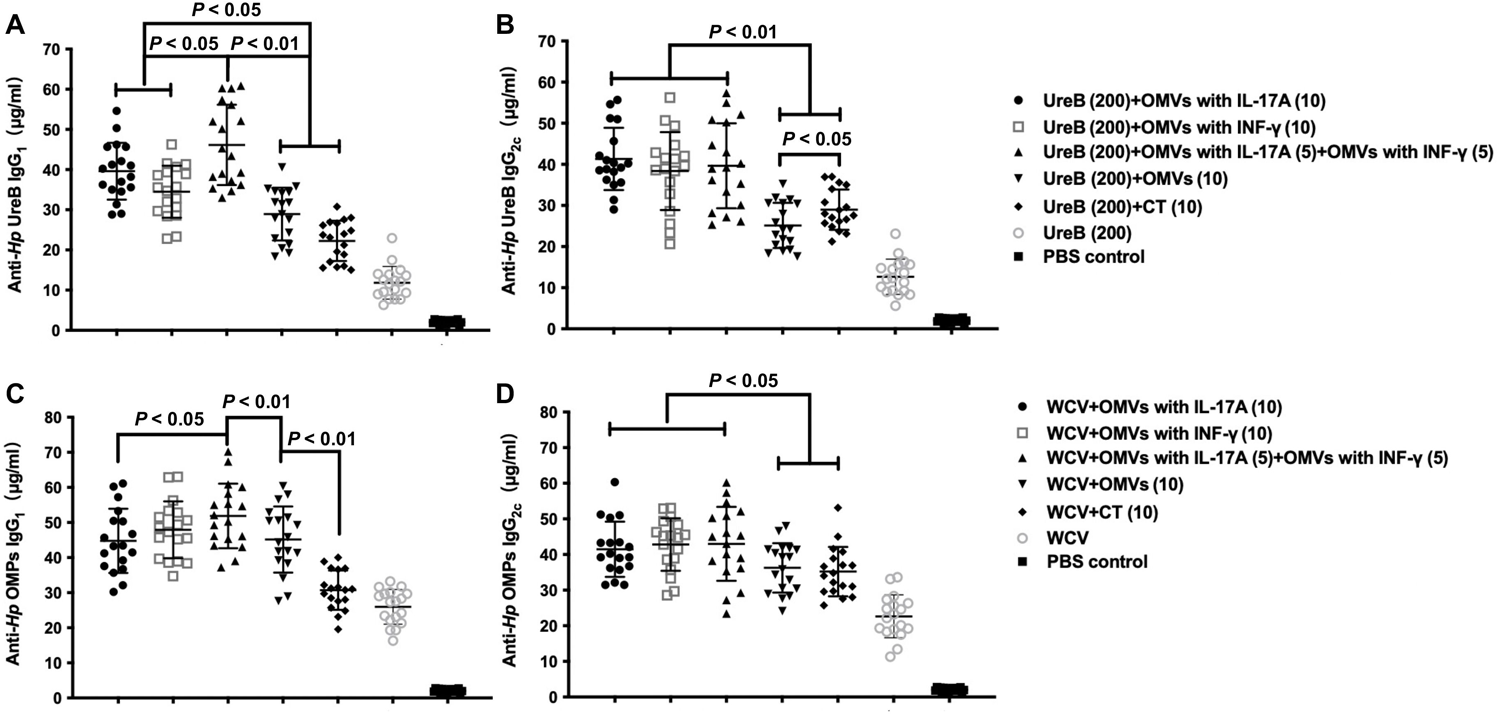
Recombinant OMVs as adjuvants induce a stronger Th1/Th17 balanced immune response. MLN cells and splenocytes of mice were isolated after 8 weeks of immunization, and the expression levels of cytokines IL-17 and IFN-γ in different groups were assessed by quantitative ELISA after immunization with UreB (A, C) or WCV (B, D) as vaccine antigens combined with recombinant OMVs or CT as adjuvants. Each group comprised nine mice, and data are expressed as mean ± SD per group. The least significant difference test was performed to determine whether the distinctions between the means of the groups were significant. *P* < 0.05 and *P* < 0.01 represent the differences between the related groups.

In addition, we evaluated the secretion of cytokines IL-4 and IL-12. We observed similar trends, with recombinant OMVs as adjuvants stimulating the production of significantly higher levels of IL-4 and IL-12 compared to the CT adjuvant (Figure 6). More interestingly, we found that the levels of IL-12 produced by recombinant OMVs as adjuvants were higher overall than IL-4 levels, regardless of whether wild-type inactivated whole-cell antigen or subunit UreB antigen was used (Figure 6). In mice, IL-12 is an indicator of a Th1 polarization response and IL-4 is an indicator of a Th2-type immune response. Thus, these data reveal that OMVs delivering cytokines as adjuvants stimulated higher levels of a Th1-type immune response. Combined with the previous data, these results suggest that recombinant OMVs as adjuvants tended to produce stronger Th1- and Th17-type immune responses. This is also consistent with previous studies, where the addition of adjuvants effectively enhanced the Th1/Th17 type immune response and thus improved the ability of the host to prevent *H. pylori* infection^22^.

**Figure 6.**
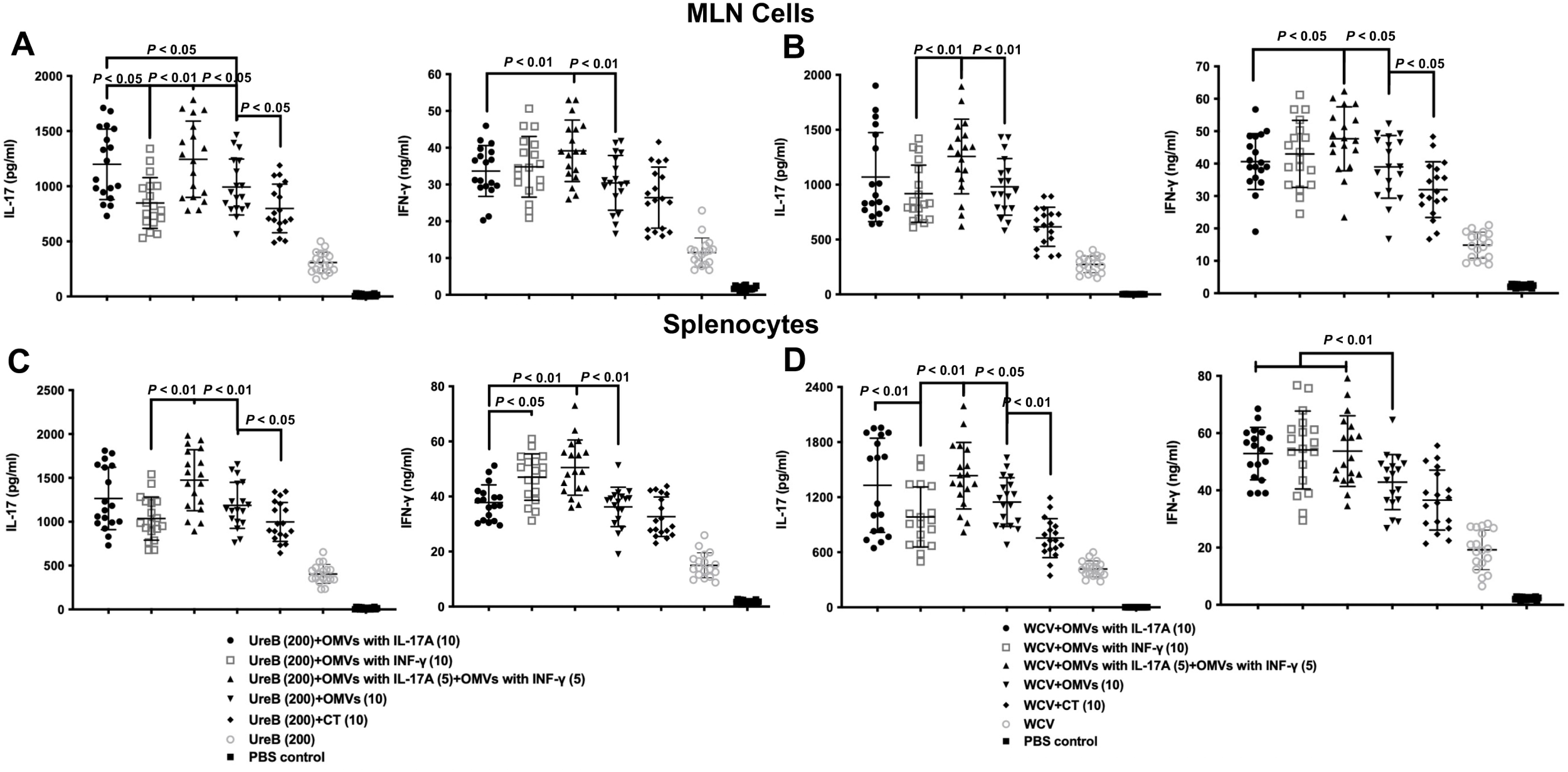
Recombinant OMVs as adjuvants induce a stronger Th1/Th2 balanced immune response. MLN cells and splenocytes of mice were isolated after 8 weeks of immunization, and the expression levels of cytokines IL-12 and IL-4 in different groups were assessed by quantitative ELISA after immunization with UreB (A, C) or WCV (B, D) as vaccine antigens combined with recombinant OMVs or CT as adjuvants. Each group comprised nine mice, and data were expressed as mean ± SD per group. The least significant difference test was performed to determine whether the distinctions between the means of groups were significant. *P* < 0.05 and *P* < 0.01 represent the differences between the related groups.

In the experiment, although the level of inflammatory factor IL-6 produced by stimulation with OMVs as adjuvants was higher than that produced by the CT adjuvant group (*P* < 0.05; Supplemental Figure 2), all experimental mice remained in good health during the immunization period and showed no abnormal behavior or death. This indicates that low doses of OMVs as adjuvants do not cause safety risks and can stimulate organismal signaling more effectively.

### 2.6 Recombinant OMVs as adjuvants protect mice from *H. pylori* infection

The mouse-adapted *H. pylori* Sydney strain 1 (SS1) challenge mouse test to evaluate vaccine protection is currently the most widely used animal model^23^. The optical density (OD) values of the urease assay were positively correlated with urease activity, reflecting the level of *H. pylori* colonization of the gastric mucosa. After immunization with either UreB or WCV with adjuvant (recombinant IFN-γ-OMVs + recombinant IL-17A-OMVs), urease activity was significantly lower (*P* < 0.01) than that in the CT adjuvant group and the wild-type *H. pylori* OMV adjuvant group (Figure 7A, 7B). We also cultured *H. pylori* isolated from gastric tissues and counted colonies to determine the density of the pathogen. The findings were consistent with the urease activity results (*P* < 0.01; Figure 7C, 7D). More interestingly, using recombinant IFN-γ-OMVs + recombinant IL-17A-OMVs as the adjuvant reduced *H. pylori* colonization in the gastric mucosa of C57BL/6 mice to a greater extent than using recombinant IL-17A-OMVs or recombinant IFN-γ-OMVs alone (*P* < 0.01; Figure 7C, 7D). Our results demonstrate that enhancing the adjuvant stimulation ability of OMVs to generate Th1/Th17-type immune responses may induce more effective immune protection against *H. pylori* infection in mice.

**Figure 7.**
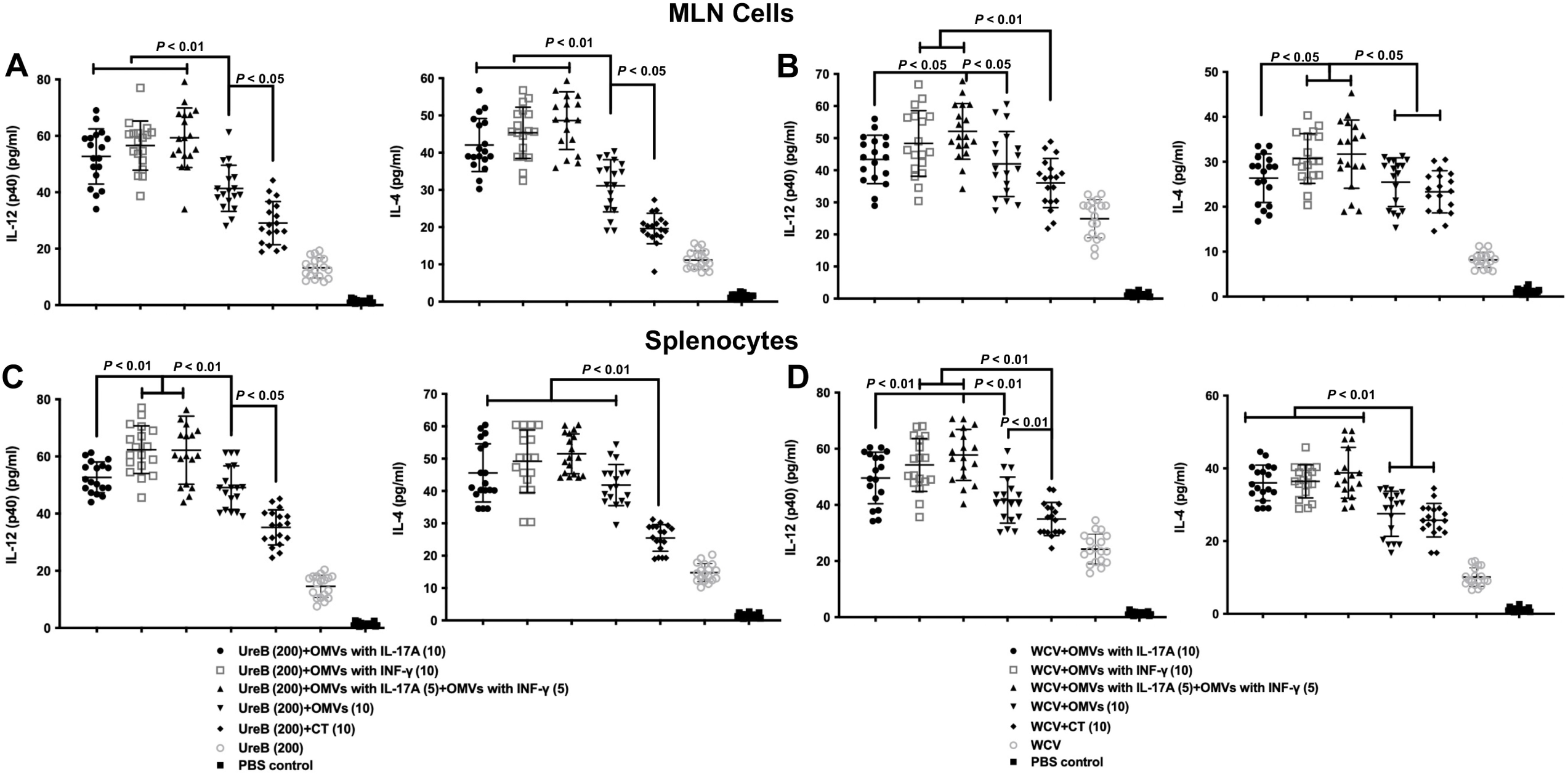
Recombinant OMVs as adjuvants elicit protection against *H. pylori* SS1 infection. Gastric tissues from immunized mice were collected and washed with PBS buffer to determine urease activity and quantify bacterial load. (A, B) Urease activity was assessed in gastric homogenates of groups of mice immunized with UreB or WCV and with recombinant OMVs or CT as adjuvants 2 weeks after the infection challenge. (C, D) *H. pylori* colony counts were quantified in the gastric prostate. Means were compared using the least significant difference test. *P* < 0.05 and *P* < 0.01 reflect statistically significant differences between relevant groups.

## 3 Discussion

In this study, we demonstrated that recombinant OMVs coupled with cytokines have significant advantages in inducing mucosal immunity or specific immune responses against two major *H. pylori* antigens compared with CT as an adjuvant, highlighting the potential applications of recombinant OMVs as adjuvants. As in previous studies, to assess the efficacy of the candidate adjuvants, we chose the *H. pylori* inactivated whole-cell vaccine and subunit vaccine UreB as antigens^24,25^, aiming to validate the immune efficacy of novel recombinant OMVs as adjuvants using these two classical vaccine forms. The adjuvant CT is widely used as a reference standard in animal models because of its ability to induce a strong mucosal response that is protective against *H. pylori* infection specifically due to the expansion of Th1 and Th17 cells in response to co-administered antigens^26^. Therefore, CT is usually chosen as the control when exploring the application potential of novel vaccine adjuvants^25^.

OMVs are non-replicating mimics of parental bacterial origin with nanoscale vesicle-like structures and contain multiple PAMPs that are ideal properties for adjuvants^27^. Our group previously explored the adjuvant efficacy of flagellin-deficient *Salmonella typhimurium* OMVs and found that flagellin-deficient OMVs stimulated stronger humoral, cellular, and mucosal immune responses in mice when co-immunized with outer membrane proteins (OMPs) as adjuvants compared to traditional aluminum adjuvants^28^. Co-inoculation of OMVs from a recombinant *Yersinia pseudotuberculosis* mutant as the adjuvant and the PcrV-HitAT fusion antigen from *Pseudomonas aeruginosa* induced an effective Th1/Th17-type immune response and rapidly cleared *Pseudomonas aeruginosa* colonization *in vivo*^29^. OMVs have been shown to exhibit adjuvant effects via multiple non-intestinal routes of administration and were superior to cholera toxin as adjuvants; additionally, even low doses of OMVs elicited adaptive immunity *in vivo*, further demonstrating the adjuvant efficacy of OMVs^30^. Of note, all the OMVs used as adjuvants in these studies were wild-type ones. While many studies have used OMVs as delivery vectors for the development of polysaccharide chimeric vaccines or tumor therapeutic vaccines, none of them have explored the potential of OMVs as delivery vectors for adjuvant development^31,32^. Here, we developed a novel strategy for adjuvants by using recombinant OMVs.

When assessing the effectiveness of a vaccine against *H. pylori* infection, the first consideration should be the activation of the mucosal immune response^26^. We examined the gastric-specific IgA antibody production stimulated by recombinant OMVs as adjuvants using quantitative ELISA and found that recombinant OMVs as adjuvants induced a stronger gastric mucosal immune response compared to the CT group (Figure 3). We also noted that the combination of recombinant OMVs with *H. pylori* vaccines significantly enhanced the level of S-IgA in vaginal secretions (Supplemental Figure 1), further demonstrating the significant effect of recombinant OMVs as adjuvants in enhancing mucosal immunity to *H. pylori* vaccines. Our data indicate that recombinant OMVs can be used as adjuvants in the development of vaccines against intestinal or vaginal pathogens.

Previous studies have shown that specific CD4+ T cells in the gastric mucosa and peripheral blood of mice infected with *H. pylori* mainly produced IFN-γ and IL-17A, two cytokines representing Th1- and Th17-type responses that are essential in inducing protective immunity against *H. pylori* and are effective in reducing the number of *H. pylori*^33,34^. However, it was not clear whether the addition of cytokines IFN-γ and IL-17A to the OMV adjuvant would be more effective. Indeed, our results demonstrated that these two cytokines could promote the adjuvant efficacy of OMVs in *H. pylori* vaccines, suggesting that the strategy of cytokine delivery by OMVs is feasible and could be used as a novel platform for adjuvant design and vaccine development.

In humans, IFN-γ, TNF-α, and IL-12 are the major cytokines involved in inducing *H. pylori*-specific Th1 cells^35^. Th2 cells primarily produce IL-4, IL-5, IL-10, and IL-13 and are responsible for producing strong antibodies that provide protection independent of phagocytes^36^. Th17 cells are the third subgroup of effector Th cells that produce IL-17, IL-17F, and IL-22, which play important roles in host defense against extracellular pathogens^37^. CT, a widely used adjuvant, has been suggested to enhance antigen-specific mucosal and humoral immunity and promote a balanced Th1/Th2/Th17 immune response^38^. In this study, we also confirmed that vaccines incorporating the adjuvant CT induced the production of Th1/Th2/Th17-type immune responses (Figures 4, 5, 6). Moreover, adjuvants based on OMVs delivering cytokines could produce higher levels of IgG_1_ and IgG_2c_ antibodies compared to the adjuvant CT (Figure 4). We also revealed that the addition of recombinant OMVs as adjuvants generated higher levels of cytokines IFN-γ, IL-17A, IL-4, and IL-12 than in the control group (Figures 5, 6). Our study suggests that recombinant OMVs as adjuvants may induce stronger Th1-, Th2-, and Th17-type immune responses. It is also important to note that recombinant IL-17A-OMVs combined with recombinant IFN-γ-OMVs as the adjuvant immunization group had lower gastric urease activity and bacterial load than the CT group and wild-type *H. pylori* OMV adjuvant group, indicating that their promotion of antigen-stimulated immune responses plays a key role in the fight against *H. pylori* infection and is more effective in eradicating *H. pylori* infection than the CT adjuvant and wild-type *H. pylori* OMV adjuvant (Figure 7).

There are some limitations in this study. First, only two candidate antigens were tested experimentally, so the compatibility of recombinant OMVs as adjuvants with other types of antigens needs to be further investigated. Second, only the adjuvant CT was selected as an experimental control to assess the adjuvant effect of recombinant OMVs in this study, a variety of commonly used *H. pylori* vaccine adjuvants should be selected for comparison to demonstrate the superiority of recombinant OMVs as adjuvants. We have clearly demonstrated that OMVs as adjuvants significantly attenuated *H. pylori* infection using urease and bacterial load assays. In future studies, we will analyze the protective efficacy of the vaccine against *H. pylori* infection in depth at the pathological level to better define the effect of recombinant OMV adjuvants.

## Conclusion

The recombinant OMVs constructed as adjuvants enhanced the production of vaccine-stimulated Th1-, Th2-, and Th17-type immune responses. In particular, the combination of recombinant IL-17A-OMVs and recombinant IFN-γ-OMVs provided stronger immune protection than CT adjuvant and wild-type *H. pylori* OMVs. Our results suggest that the construction of novel *H. pylori* vaccines using OMVs as adjuvants to deliver cytokines that effectively enhance Th1/Th2/Th17 immune responses may represent a promising solution for the development of more effective *H. pylori* vaccines.

## 4 Materials and methods

### 4.1 Bacterial culture and OMV preparation

The gerbil-adapted *H. pylori* 7.13 strain derived from clinical strain B128 was kindly provided by Dr. Yong Xie at the First Affiliated Hospital of Nanchang University in Nanchang, China. *H. pylori* SS1 suspensions used for challenging were prepared from fresh exponential-phase cultures to ensure a high number of viable cells. All *H. pylori* strains in this study were cultured in Campylobacter agar base supplemented with 10% sheep blood (Difco, Detroit, MI, USA) at 37 °C with 5% O_2_, 10% CO_2_, and 85% N_2_.

OMVs from the *H. pylori* 7.13 strain were isolated via ultracentrifugation as previously described^39^. Briefly, 500 mL of the *H. pylori* 7.13 strain was cultured to the logarithmic phase (OD at 600 nm [OD600] of 1; 48 to 72 h) and centrifuged at 4,500 *g* and 4 °C for 1 h to remove bacterial particles. Then, the supernatant was filtered twice with the cap of a 0.45 mm sterilized filter (Millipore, Billerica, MA, USA). The filtrate containing *H. pylori* OMVs was ultracentrifuged at 20,000 *g* and 4 °C for 2 h, and the OMV particles were washed with phosphate buffered saline (PBS) (Mediatech, Manassas, VA, USA). The pelleted OMV particles were resuspended in OptiPrep density gradient medium (Sigma-Aldrich, St. Louis, MO, USA) in PBS and ultracentrifuged for 24 h at 100,000 *g* and 4 °C in a density gradient of 40%, 35%, 30%, and 20%. The OMV fractions were pooled, gently washed three times with PBS, dissolved in 1 mL PBS, and stored at −20 °C. For antigen coating in the ELISA and antigen stimulation in cytokine detection, OMPs were isolated from the *H. pylori* 7.13 strain as previously described^40^. The protein concentrations of the obtained OMVs and OMPs were quantified using a bicinchoninic acid (BCA) assay kit (Thermo Fisher, Rockford, IL, USA) according to the manufacturer’s instructions.

### 4.2 Cell culture

The human embryonic kidney cell line HEK-293T and human gastric mucosal epithelial cells GES-1 were purchased from the American Type Culture Collection (ATCC, Manassas, VA, USA). Briefly, HEK-293T cells were cultured in Dulbecco’s Modified Eagle Medium (DMEM)/F12 medium (Cat# D8437, Sigma, MO, USA) supplemented with 10% fetal bovine serum (FBS; Cat# 10091-148, Gibco, CA, USA), streptomycin (100 mg/ml), and penicillin (100 U/ml) at 37 °C with 5% CO_2_.

### 4.3 Laser scanning confocal microscope analysis

After co-incubation of OMVs delivering cytokines with cells, the expression of fluorescent proteins EGFP and mCherry in the cells was observed using a laser scanning confocal microscope (OLYMPUS, Tokyo, Japan), and representative images were obtained.

### 4.4 ELISA quantification of cytokines IFN-γ and IL-17A

After co-incubation of OMVs delivering cytokines with cells for 24 h, the cells were collected and lysed, the supernatant was collected by centrifugation, and the concentrations of IFN-γ and IL-17A in the supernatant were determined by standard ELISA.

### 4.5 RT-qPCR

TRIzol reagent (Invitrogen, 15596018) was used to extract total RNA, and HiScript III RT SuperMix (Vazyme, R323-01) was employed to perform the reverse transcription reaction. Quantitative RT-PCR (qPCR) experiments were conducted with ChamQ SYBR Color qPCR Master Mix (Vazyme, Q431-02). Briefly, the cycle program of qPCR consisted of denaturation (95 °C, 15 s), annealing (60 °C, 15 s), and extension (72 °C, 45 s). Raw data were processed with the 2–ΔΔCt method.

### 4.6 Animal experiments

All animal experiments were approved by the Animal Welfare Committee of Nanchang University (Nanchang, China; approval number NCDXYD-202102), and followed the relevant rules and the guidelines for the care and use of laboratory animals. We made every effort to minimize animal pain throughout our experiments. Female C57BL/6 mice (6 weeks old, 16–22 g) were purchased from the Experimental Animal Science Center of Nanchang University. After 1 week of adaptation to the new environment, mice were divided into 14 groups. Each group contained nine mice that received the first vaccine injection by gavage (0 days). The immunogens selected and the doses for each group are described in Table 1. UreB and WCV were used as vaccine antigens, and *H. pylori* strain WCV consisted of 10^9^ inactivated cells. UreB, CT (Sigma-Aldrich, St. Louis, MO, USA), and OMV were suspended in 200 μL PBS buffer. The PBS alone (10 μL) group served as a negative control. Blood samples were collected by orbital sinus puncture the day before the first immunization and 2, 4, 6, 8, 10, and 12 weeks later; vaginal secretions were collected using five flushes of 0.1 mL PBS. Subsequently, the soluble fraction of serum and vaginal secretions was obtained by centrifugation. Intragastric inoculation with the appropriate antigen was performed in week 4 for intensive immunization. Splenocytes were collected for cytokine assay in week 8 at the expense of some mice. In week 14 after initial immunization, all remaining mice were challenged with 10^9^ colony-forming units (CFU) of *H. pylori* SS1 in 20 μL of PBS containing 0.01% gelatin by the oral route and continuously monitored until week 16. Finally, all mice were sacrificed and their stomach tissues were harvested for urease assay and bacterial load determination. The immunization timeline is shown in Figure 2A. All animal experiments were repeated twice, and data were finally collected for statistical analysis.

**Table 1.**
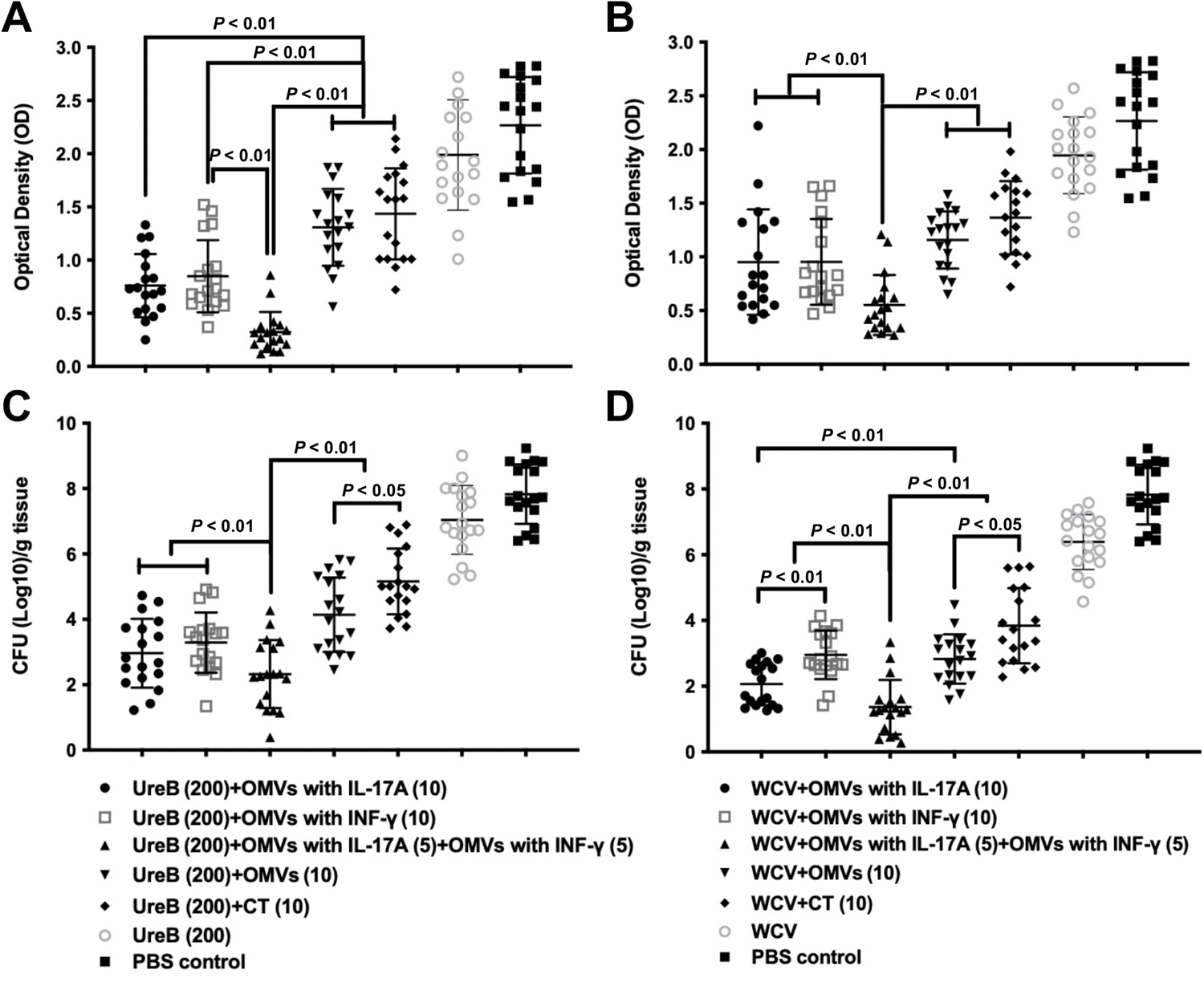
Vaccine formulation strategy for immunization using recombinant *H. pylori* OMVs as adjuvants*.

### 4.7 ELISA

Antibody levels in mouse blood samples and vaginal secretions were quantified using quantitative ELISA as described previously^41^. OMP (1 mg) suspended in 100 mL of sodium carbonate buffer (pH 9.6) was used to coat each well of a 96-well plate (Nalge Nunc Inc., Naperville, IL, USA) that was incubated overnight at 4 °C. Purified mouse Ig isotype standards (IgG, IgG_1_, IgG_2c_, and IgA; BD Biosciences, Billerica, MA, USA) were prepared in triplicate and diluted twice (0.5 mg/mL). After three washes with PBS containing 0.1% Tween 20 (PBST), the plate was blocked with 2% bovine serum albumin (BSA) solution for 2 h at room temperature. Subsequently, 100 mL of each sample was added to the respective wells at different dilutions in triplicate, and the plate was incubated at room temperature for 1 h. After three washes with PBST, biotinylated goat anti-mouse antibodies IgG, IgG_1_, IgG_2c_, and IgA (Southern Biotechnology Inc., Birmingham, AL, USA) were added to each well. Then, streptococcal protease alkaline phosphatase conjugate (Southern Biotechnology) was added, and the substrate p-nitrophenyl phosphatase (Sigma-Aldrich) in diethanolamine buffer (pH 9.8) was used to develop the wells. Absorbance was measured at 405 nm on an automatic ELISA plate reader (model EL311SX; Biotek, Winooski, VT, USA). The final Ig isotype concentration of antibody samples was calculated separately for each antibody isotype using a standard curve.

### 4.8 Detection of cytokines in mouse MLN cells and splenocytes

Mouse MLN cells and splenocytes were collected after 4 weeks of booster immunization and stimulated with 6 mg/mL of OMP isolated from *H. pylori* strain 7.13 for 24 h as previously described^18^. Supernatants from stimulated MLN cells and splenocytes were then collected and cytokine levels were quantified by ELISA. Monoclonal antibodies against IFN-γ, IL-17A, IL-12 (p40), IL-4, and IL-6 were added to 96-well plates (BD Biosciences, Mountain View, CA, USA). Next, samples were blocked with PBS containing 1% BSA, added to the wells in triplicate, and incubated overnight at 4 °C. Then, the wells were washed and incubated with biotinylated monoclonal anti-IFN-γ, anti-IL-4, anti-IL-17A, anti-IL-12 (p40), and anti-IL-6 antibodies (BD Biosciences, Billerica, MA, United States). Finally, horseradish peroxidase-labeled anti-biotin antibody (Vector Laboratories, Burlingame, CA, USA) was added to each well along with 3,39,5,59-tetramethylbenzidine (TMB; Moss Inc., Pasadena, CA, USA) to enhance the reaction, which was terminated with 0.5 M hydrochloric acid (HCl). Standard curves were created based on mouse recombinant (r) IFN-γ, IL-17A, IL-4, IL-12 (p40), and IL-6 to determine cytokine expression levels in spleen cells.

### 4.9 Determination of bacterial loading

Two weeks after the oral *H. pylori* SS1 challenge, gastric tissues were collected to quantify bacterial load. A lower pathogen load indicates a stronger protective effect. First, the separated tissue was washed with precooled PBS and then transferred to pre-weighed test tubes containing 5 mL of brain heart infusion medium (Difco, Detroit, Mich., USA). Next, the tissue was reweighed to an accuracy of 0.0001 g, homogenized with a sterile homogenizer, and continuously inoculated onto a Campylobacter agar matrix (Difco, Detroit, Mich., USA) plate containing 10% sheep blood at dilutions of 1:10, 1:100, and 1:1,000. Then, the plates were incubated at 37 °C under microaerobic conditions for 6 to 7 days. *H. pylori* colonies were identified based on the urease and oxidase reactions and wet morphology analysis.

### 4.10 Urease test

Stomach specimens from each mouse were immersed in 0.5 mL of 0.8% sodium chloride (NaCl) solution to prepare a tissue homogenate for urease activity quantification^42^. Briefly, 3 mL of urea broth (1 mg/mL glucose, 1 mg/mL peptone, 2 mg/mL monopotassium phosphate (KH_2_PO_4_), 5 mg/mL NaCl, and 1% urea) containing phenol red indicator was mixed with 100 mL of the tissue mixture and homogenized. Tissue homogenate containing PBS was used as the negative control. After incubating at 37 °C for 4 h, the urease activity of each gastric tissue sample was quantified based on the OD550 using a UV/visible spectrophotometer.

### 4.11 Statistical analysis

All ELISA experiments were performed in triplicate. The significance of differences in the average values between the experimental and control groups was assessed using one- or two-way analysis of variance (ANOVA), followed by Tukey’s post hoc test. All data are expressed as mean ± standard deviation (SD). All statistical analyses were performed using GraphPad Prism software v.9.0 (GraphPad Software Inc., San Diego, CA, USA).

## Acknowledgments

This study was supported by the National Natural Science Foundation of China (82203032, 32260193), Training Plan for Academic and Technical Leaders of Major Disciplines in Jiangxi Province-Youth Talent Project (20212BCJ23036), Project for high and talent of Science and Technology Innovation in Jiangxi “double thousand plan” (S2021GDQN1008) and the start-up fund from the First Affiliated Hospital of Nanchang University

## Declarations

### Author contributions

Qiong Liu, Tian Gong and Chengsheng Zhang conceived and designed the experiments; Biaoxian Li, Jiahui Lu, Yejia Zhang, Yinpan Shang, and Yi Li performed the experiments. Biaoxian Li, Qiong Liu, Tian Gong and Chengsheng Zhang analyzed the data; Biaoxian Li, Qiong Liu and Chengsheng Zhang wrote the manuscript.

### Data availability statements

The datasets generated during and analysed during the current study are available from the corresponding author on reasonable request.

---

**Supplemental Figure 1. Effect of recombinant OMVs as adjuvants on mucosal immune responses to different types of antigens.** Quantitative ELISA was performed to determine the level of anti-*Hp* UreB in the secretions of each group within 12 weeks of immunization when UreB was used as an antigen (A) and the level of anti-*Hp* OMP when WCV was used as an antigen (B). Statistical significance was assessed by two-way ANOVA. *P* < 0.05 was considered statistically significant. All the results are expressed as mean ± SD per cohort.

**Supplemental Figure 2. Safety assessment of recombinant OMVs as adjuvants.** After 8 weeks of immunization, mouse MLN cells and splenocytes were isolated and the expression levels of cytokine IL-6 in different groups were assessed by quantitative ELISA after immunization with UreB (A, C) or WCV (B, D) as vaccine antigens combined with recombinant OMVs or CT as adjuvants. Each group consisted of nine mice, and data were expressed as mean ± SD. The least significant difference test was performed to determine the significance of the difference between the means of the groups. *P* < 0.05 represent the differences between the groups of interest.

